# Communicating compositional patterns

**DOI:** 10.1101/451161

**Authors:** Eric Schulz, Francisco Quiroga, Samuel J. Gershman

## Abstract

How do people perceive and communicate structure? We investigate this question by letting participants play a communication game, where one player describes a pattern, and another player redraws it based on the description alone. We use this paradigm to compare two models of pattern description, one compositional (complex structures built out of simpler ones) and one non-compositional. We find that compositional patterns are communicated more effectively than non-compositional patterns, that a compositional model of pattern description predicts which patterns are harder to describe, and that this model can be used to evaluate participants’ drawings, producing human-like quality ratings. Our results suggest that natural language can tap into a compositionally structured pattern description language.

## Introduction

Humans see patterns everywhere, and eagerly communicate them to one another. However, little is known formally about how we communicate patterns, what kinds of patterns are easier or harder to communicate, and how we reconstruct patterns from natural language. This paper seeks to bridge this gap by combining a pattern communication game with a mathematical model of pattern description (Quiroga, Schulz, Speekenbrink, & Harvey, 2018; Schulz, Tenenbaum, Duvenaud, Speekenbrink, & Gershman, 2017).

Consider the graphs shown in Figure 1, which plot time series of CO2 emission, airline passenger volume, search frequency for the term “gym membership.” Experiments suggest that humans perceive these graphs as compositions of simpler patterns, such as lines, oscillations, and smoothly changing curves (Quiroga et al., 2018; Schulz, Tenenbaum, et al., 2017). For example, there is seasonal variation in passenger volume (a periodic component with time-dependent amplitude), superimposed on a linear increase over time.

**Figure 1.**
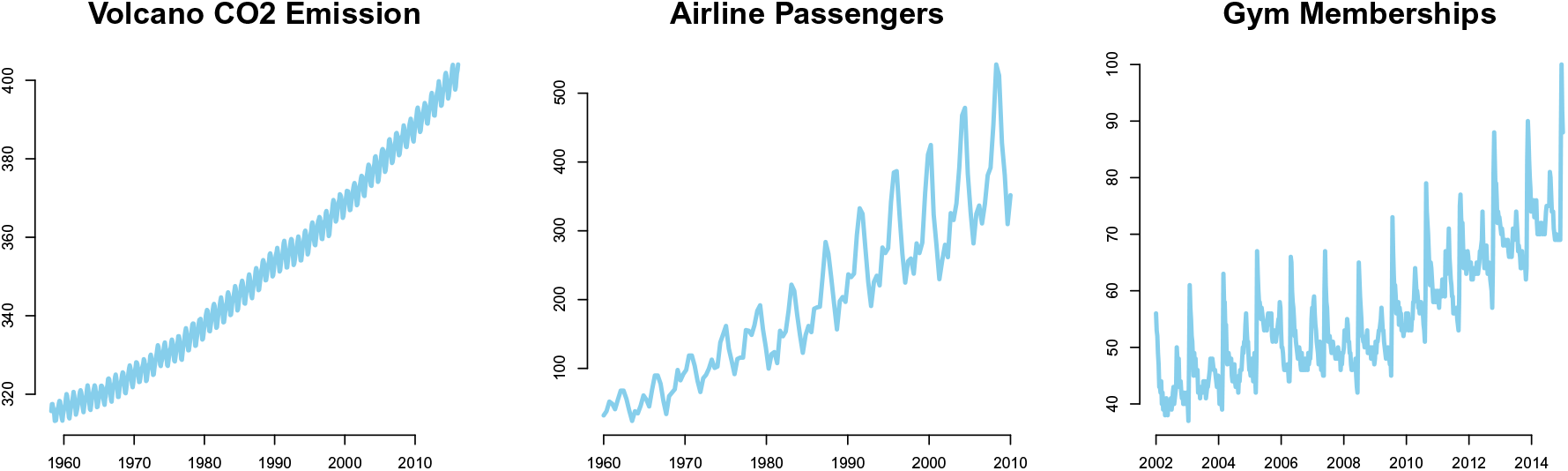
Examples of compositional patterns. Left: Monthly average atmospheric CO2 concentrations collected at the Mauna Loa Observatory in Hawaii from 1960-2010. Center: Number of airline passengers from 1960-2010, originally collected by Box, Jenkins, Reinsel, and Ljung (2015). Right: Google queries for “Gym membership” from 2002-2012 in the city of London.

As described in more detail in the next section, we can formalize this idea using a pattern description language consisting of functional primitives and algebraic operations that compose them together. By defining a probability distribution over this description language, we can express an inductive bias for certain kinds of functions—in particular, functions that can be described with a small number of compositions (Duvenaud, Lloyd, Grosse, Tenenbaum, & Ghahramani, 2013; Lloyd, Duvenaud, Grosse, Tenenbaum, & Ghahramani, 2014; Schulz, Tenenbaum, et al., 2017). In other words, the “mental” description length of a function relates to the complexity of its encoding in the compositional pattern description language.

It is important to note that there are other ways to reduce description length besides encoding functions with a small set of compositions (what we will refer to as “compositional functions”). For example, a standard assumption in machine learning is that functions are smooth (Rasmussen & Williams, 2006). If we defined a probability distribution over functions that prefers smoothness, then smooth functions would have short description lengths, in the sense that the number of bits required to encode them would be smaller than non-smooth functions. However, a preference for smoothness does not seem to be an adequate account of how humans encode functions: functions that are smooth but cannot be compactly described by compositions are less easily encoded, as indicated by poorer memory and change detection performance for these functions compared to compositional functions (Schulz, Tenenbaum, et al., 2017, see also additional analysis in the Supporting Information).

Here we extend this idea one step further, asking whether there is a correspondence between the pattern description language and natural language descriptions of functions. We proceed in three steps. First, we ask participants to describe functions sampled from compositional or non-compositional distributions. Second, we asked a separate group of participants to redraw the original function using only the description. Third, we ask another group of participants to rate how well each drawing corresponds to the original. We hypothesized that compositional functions would be easier to reconstruct compared to non-compositional functions, under the assumption that the former allow for a mental description that can be more easily encoded into natural language and decoded back into the function space. We also rule out several alternative explanations and map pattern-specific descriptions to compositional components with the help of an additional experiment.

## A compositional pattern description language

Our model of pattern description is based on a Gaussian Process (GP) regression approach to function learning (Rasmussen & Williams, 2006; Schulz, Speekenbrink, & Krause, 2017). A GP is a collection of random variables, any finite subset of which is jointly Gaussian. A GP defines a distribution over functions. Let *f* : *X* → ℝ denote a function over an input space *X* that maps to real-valued scalar outputs. This function can be modeled as a random draw from a GP:

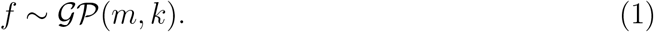

The mean function *m* specifies the expected output of the function given input **x**, and the kernel function *k* specifies the covariance between outputs.

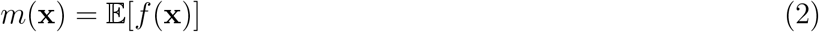

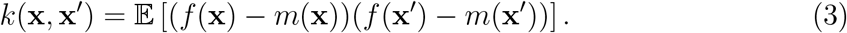

We follow standard convention in assuming a prior mean of **0** (Rasmussen & Williams, 2006).

All positive semi-definite kernels are closed under addition and multiplication, allowing us to create richly structured and interpretable kernels from well-understood base components. We use this property to construct a class of compositional kernels (Duvenaud et al., 2013; Lloyd et al., 2014; Schulz, Tenenbaum, et al., 2017). To give some intuition for this approach, consider again the C02 data in Figure 1. This function is naturally decomposed into a sum of linearly increasing component and a seasonally periodic component. The compositional kernel captures this structure by summing a linear and periodic kernel.

Compositional GPs have been used to model complex time series data (Duvenaud et al., 2013), as well as to generate automated natural language descriptions from data (Lloyd et al., 2014), an approach coined the “automated statistician” (Ghahramani, 2015). Although it is frequently assumed that people will easily understand the generated description of the “automated statistician”, it is not known whether compositional patterns are indeed more communicable.

We follow the approach developed in Schulz, Tenenbaum, et al. (2017), using three base kernels that define basic structural patterns: a linear kernel that can encode trends, a radial basis function kernel that can encode smooth functions, and a periodic kernel that can encode repeated patterns (see Tab. 1). These kernels can be combined by either multiplying or adding them together. In previous research, we found that this compositional grammar can account for participants’ behavior across a variety of experimental paradigms, including pattern completions, change detection, and working memory tasks (Schulz, Tenenbaum, et al., 2017). We fix the maximum number of combined kernels to be three and do not allow for repetition of kernels in order to restrict the complexity of inference (see next section).

**Table 1.**
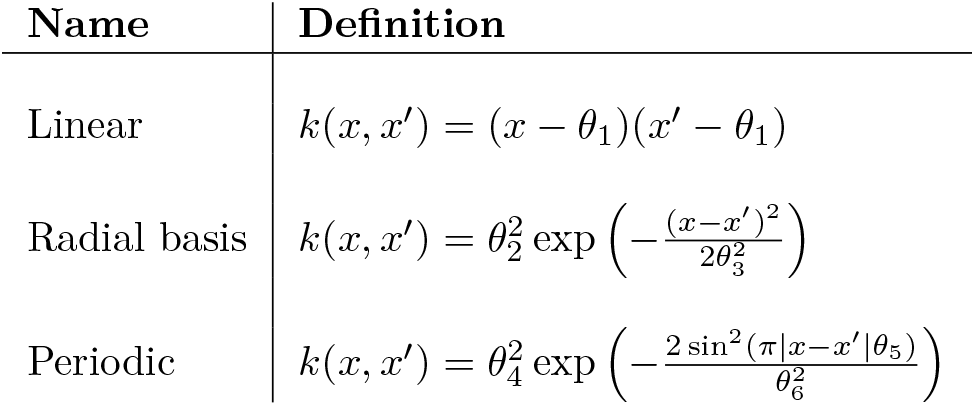
Base kernels in the compositional grammar.

We compare the compositional model to a non-compositional model based on a spectral mixture of kernels (see Supporting Information for further details). This model is derived from the fact that any stationary kernel can be expressed as an integral using Bochner’s theorem. This model approximates functions by matching their spectral density with a mixture of Gaussians. It has a similar expressivity compared to the compositional model, but does not encode compositional structure explicitly. This means that both models will make similar predictions given unlimited data; however, given a finite data regime the compositional kernel will have strong inductive biases for compositional functions, whereas the spectral kernel will not show such inductive biases. Wilson, Dann, Lucas, and Xing (2015) have used this model to reverse-engineer “human kernels” in standard function learning tasks. We use this kernel to asses if communication of patterns can be described well by a kernel that is equally expressive as the compositional kernel but does not operate over structural building blocks. Instead of optimizing its parameters to find human-like kernels in traditional function learning tasks, we will optimize it based on the structure participants had to describe^1^.

## Modeling function learning

We model human pattern description using a Bayesian inference over functions with a GP prior, an approach that has been successfully applied to a range of experimental and observational data (Griffiths, Lucas, Williams, & Kalish, 2009; Lucas, Griffiths, Williams, & Kalish, 2015; Schulz et al., 2019; Wu, Schulz, Speekenbrink, Nelson, & Meder, 2018). Given an observed pattern 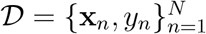, where *y_n_ ∼ N* (*f* (**x***_n_*)*, σ*^2^) is a draw from the latent function, the posterior predictive distribution for a new input **x***_*_* is also normally distributed, where

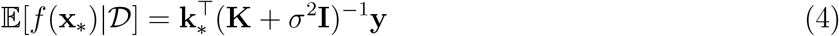

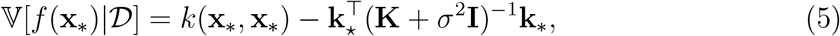

are the mean and variance respectively. The term **y** = [*y*_1_*, …, y_N_*]*^T^*, **K** is the *N × N* matrix of covariances evaluated at each pair of observed inputs, and **k***_*_* = [*k*(**x**_1_, **x***_*_*)*, …, k*(**x***_N_*, **x***_*_*)] is the covariance between each observed input and the new input **x***_*_*.

We use a Bayesian model comparison approach to evaluate how well a particular kernel captures the data, while accounting for model complexity. Assuming a uniform prior over kernels, the posterior probability favoring a particular kernel is proportional to the marginal likelihood of the data under that model. The log marginal likelihood for a GP with hyper-parameters *θ* is given by:

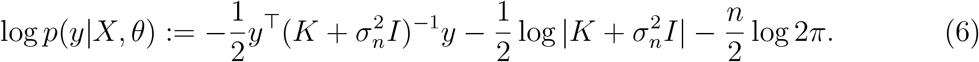

where the dependence of *K* on *θ* is left implicit. The hyper-parameters are chosen to maximize the log-marginal likelihood, using gradient-based optimization (Rasmussen & Nickisch, 2010).

## Generating patterns

We use the same patterns as in Schulz, Tenenbaum, et al. (2017). These patterns were generated from both compositional and non-compositional (spectral mixture) kernels. The compositional patterns were sampled randomly from a compositional grammar by first randomly sampling a kernel composition and then sampling a function from that kernel, whereas the non-compositional patterns were sampled from the spectral mixture kernel, where the number of components was varied between 2 and 6 uniformly. A subset of these sampled patterns were then chosen so that compositional and non-compositional functions were matched based on their spectral entropy and wavelet distance (Goerg, 2013), leading to a final set of 40 patterns.

## Pattern communication game

Our study assessed how well different patterns can be communicated in a free form communication game (i.e., without any restrictions on participants’ description lengths or word usage). The study consisted of three parts: description, drawing, and quality rating. Participants were recruited from Amazon Mechanical Turk, and no participant was allowed to participate in more than one part. The study was approved by Harvard ethic’s review board.

### Part 1: Eliciting descriptions

31 participants (6 female, mean age=34.91, SD=10.25) took part in the description study. Participants sequentially saw 6 different patterns, represented as graphs which they had to describe afterwards. Three of the patterns were randomly sampled from the 20 compositional patterns without replacement, and three were sampled from the non-compositional pool of patterns. The order of the presented patterns was determined at random. On every trial, participants first saw a pattern for 10 seconds, after which the pattern disappeared. The pattern was shown to them as 100 equidistant points indicating a function on a canvas (see Fig. 3). After the pattern disappeared, participants had to describe it using as many words as they liked. Participants were told that we would pass on their descriptions to someone else who would then have to redraw the patterns without ever having seen them.

**Figure 2.**
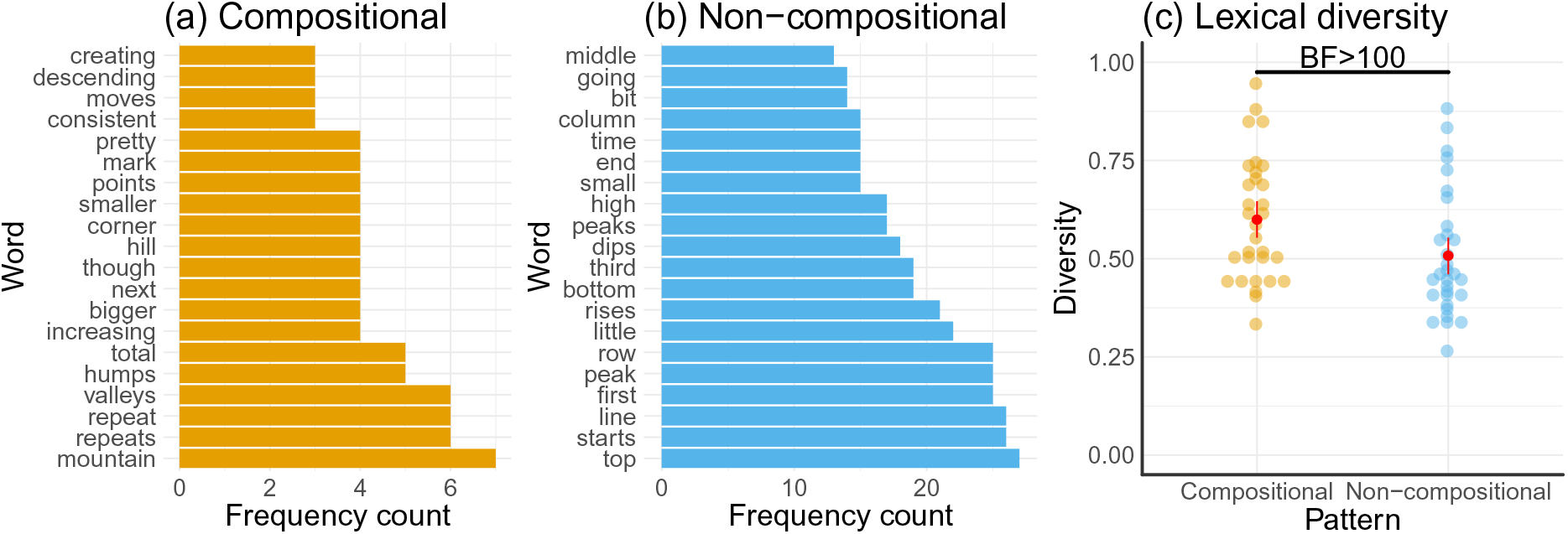
**a:** Frequency of words that were used more than twice in the compositional but not the non-compositional descriptions. **b:** Frequency of words that were used more than twice in the non-compositional but not the compositional descriptions. c: Lexical diversity of compositional and non-compositional descriptions.

**Figure 3.**
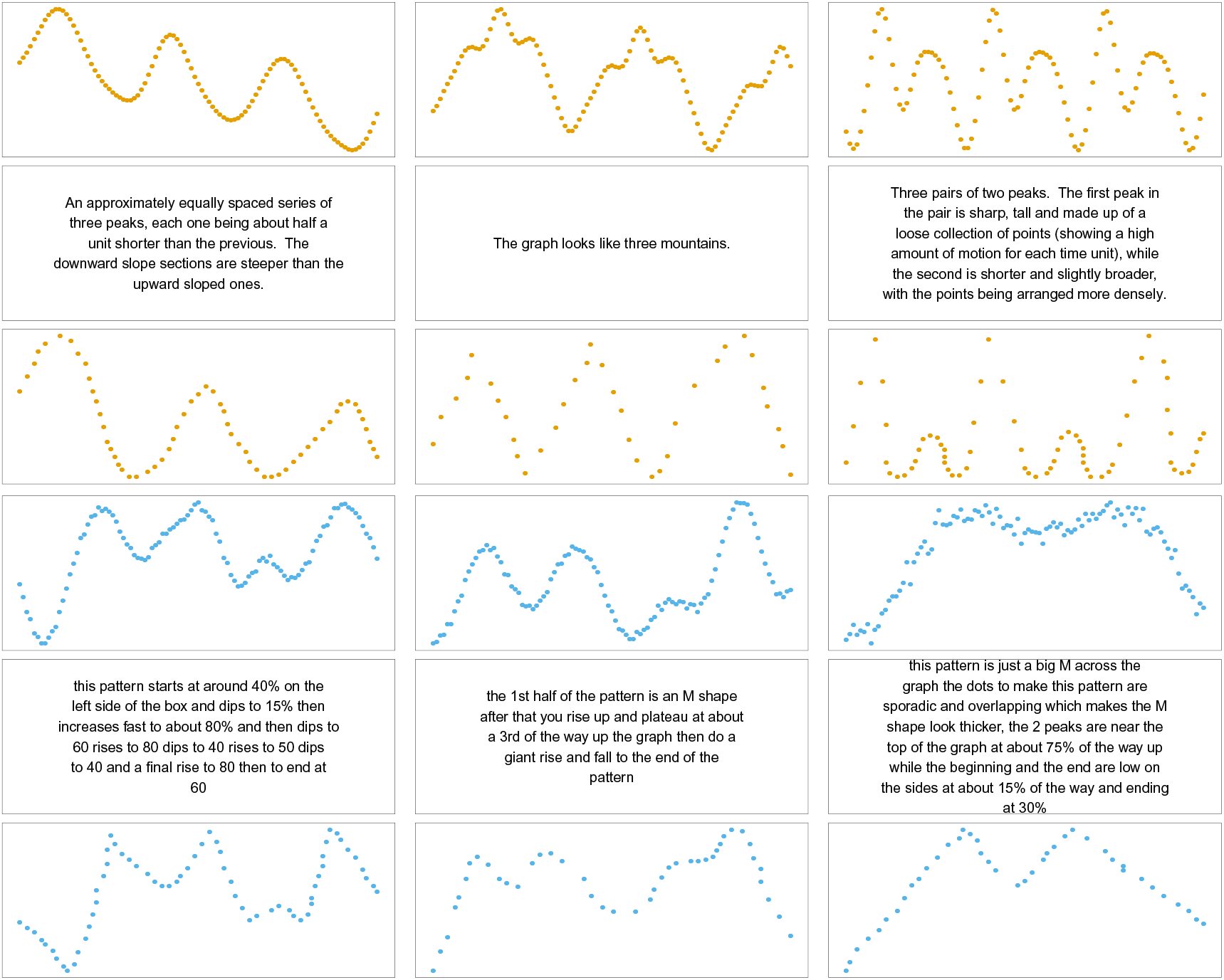
Examples of descriptions and drawings. Figures show the 3 best (based on the quality ratings) unique drawings for both compositional (upper panel in orange) and non-compositional (lower panel in blue) patterns. The upper rows always show the original pattern, the middle rows show the descriptions, and the bottom rows show the redrawn patterns.

Two judges independently rated the descriptions^2^ on a scale from 1 (bad descriptions) to 5 (great descriptions). The agreement between the two judges was sufficiently high, with a inter-rater correlation of *r*(29) = 0.46, *t* = 2.45, *p* = .02, *BF* = 3.8, and we validated their judgements both statistically and using additional raters (see Supporting Information). We then retained the descriptions with average rating higher than 3, giving 7 “describers” and a total pool of 31 different patterns. Sixteen of these patterns were compositional, and fifteen were non-compositional. All participants were paid $2 for their participation.

### Part 2: Drawing the patterns

We recruited 49 participants (21 females, mean age=33.6, SD=9.6) for the drawing part of the experiment. In this part, participants only saw the descriptions of the patterns and had to redraw them by placing dots on an empty canvas. Below the canvas, participants saw the descriptions of the patterns, which they knew had been written by a past participant. Participants were told that they could place any number of dots onto the canvas, but had to place at least 5 dots to draw a pattern before they could submit their drawings. Each participant received the 6 descriptions written by a randomly-matched participant from the description part, i.e. they were paired with one of the top 7 “describers” from the first part of the study. Participants were paid $2 for their participation.

### Part 3: Rating the quality of the drawings

104 participants (35 females, mean age= 37.7, SD=8.6) were recruited to rate the quality of participants’ performance in the previous parts. Participants were told the rules of the game the previous participants had played. They then had to rate 30 randomly sampled drawings, where the drawings were always presented right next to the original pattern. Participants did not see the descriptions that lead to the eventual drawings, but rather only had to evaluate how much the drawing resembled the original, i.e. how well they thought two participants performed in one round of the game. They did this by entering values on a slider from 0 (bad performance) to 100 (great performance). We paid participants $1 for their participation.

## Results

Figure 3 shows three examples of participants’ descriptions and drawings for both compositional and non-compositional patterns. We first assessed whether participants in the description part of the study entered longer descriptions for the compositional than the non-compositional patterns. This analysis revealed no significant difference between the two kinds of patterns (*t*(30) = 0.15, *p* = .88, *d* = 0.03. *BF* = 0.2). Next, we assessed whether participants in the drawing part of the study used more dots to redraw compositional than non-compositional patterns. This also showed no difference between the two kinds of patterns (*t*(49) = 1.00, *p* = .32, *d* = 0.14, *BF* = 0.2).

Although one might conclude from these analyses that the descriptions and redrawings were relatively similar across the two pattern classes, inspection of which words frequently appeared in the compositional descriptions but not the non-compositional descriptions (and vice versa) revealed that compositional descriptions often included more abstract words such as “mountain”, “repeat” or “valley” (Fig. 2a), whereas non-compositional descriptions used words such as “starts”, “bottom” or “top”, likely describing exactly how to draw a particular shape (Fig. 2b). Furthermore, we assessed the descriptions’ lexical diversity, defined as the sum of the unique words used divided by all words used in a description (McCarthy & Jarvis, 2010). Compositional descriptions showed a higher lexical diversity than non-compositional descriptions (*t*(30) = 4.22, *p* < .001, *d* = 0.76, *BF >* 100, Fig. 2c).

We next analyzed the quality of participants’ drawings. In order to compare the two, we used polynomial smoothing splines to connect the dots. The splines were forced to go through every point on the canvas such that the original and redrawn patterns have the same length. Our results also hold even if we just use the raw points or other methods of extracting the patterns such as generalized additive models (see Supporting Information). We then calculated the absolute difference (absolute error) between the original and the redrawn patterns. This difference was larger for non-compositional than for compositional patterns (Fig. 4a; *t*(49) = 2.43, *p* = .01, *d* = 0.34, *BF* = 4.1), indicating that participants were more accurate at redrawing compositional patterns.

**Figure 4.**
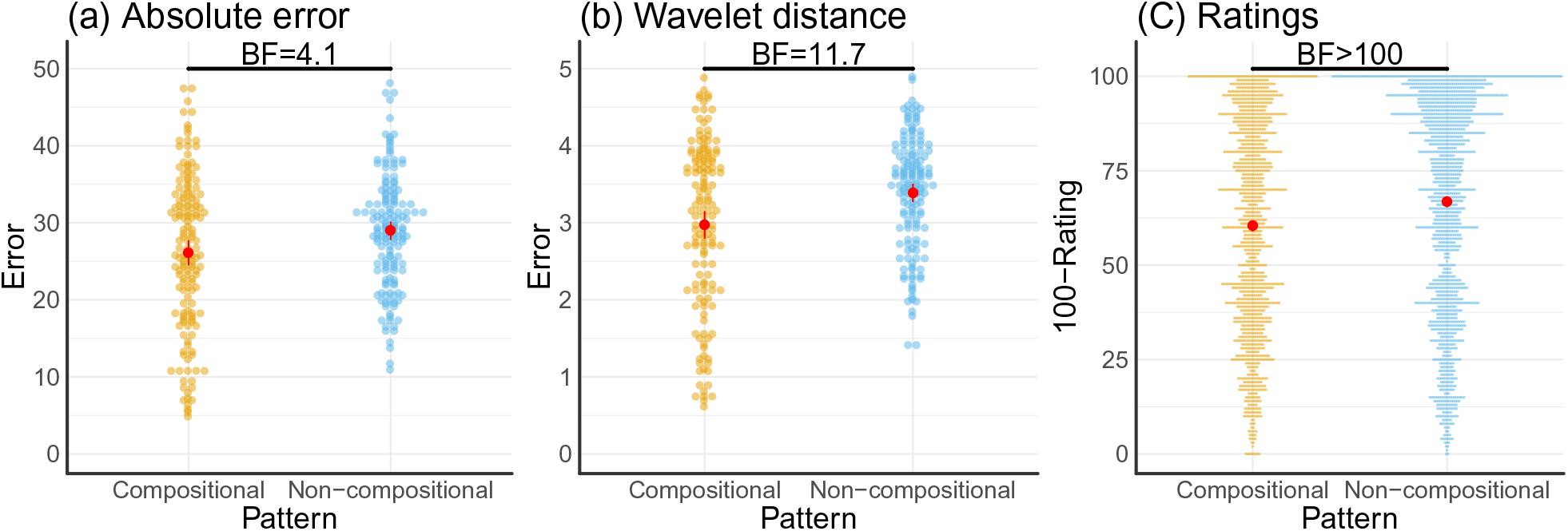
Difference between compositional and non-compositional functions. Colors indicate the type of pattern. Red dots show the mean, along with the 95% confidence interval. a: Absolute error between original and redrawn patterns. b: Wavelet distance between original and redrawn patterns. c: Rated quality shown as 100-Rating to transform it to a distance measure (i.e. lower values are better).

The absolute distance between two patterns might not be the best indicator of performance, because two patterns can look alike but still show a large absolute difference (e.g., if the redrawn pattern is smaller than the original, or if one pattern is just slightly shifted to either side). We therefore also applied a distance measure that takes into account these possible deviations by assessing the similarity of two patterns based on their differences after performing a Haar wavelet transform. The idea behind this similarity measure is to replace the original pattern by its wavelet approximation coefficients, and then to measure similarity between these coefficients (see Supporting Information, Montero, Vilar, et al., 2014). Technicalities aside, this measure is robust to scaling and shifting of the patterns. We have previously verified that it corresponds well with participants’ similarity judgments when comparing two patterns (Schulz, Tenenbaum, et al., 2017). Analyzing participants’ performance using this measurement (Wavelet distance) showed an even stronger advantage for compositional patterns (Fig. 4b; *t*(49) = 3.02, *p* = .004, *d* = 0.43, *BF* = 11.7).

Next, we looked at the quality ratings collected in the third part of our study. We estimated a linear mixed-effects model with random effects for a compositional vs. non-compositional contrast for raters, describer-drawer pairs, and for the items (patterns). We compared this model to another model that also included a compositional vs. non-compositional contrast as a fixed effect (following the logic of Barr, Levy, Scheepers, & Tily, 2013). The results of this analysis showed that adding the compositional contrast as a fixed effect moderately improved the overall model fit (*BF* = 4.6). Compositional patterns were rated more highly than non-compositional patterns (Fig. 4c), resulting in a posterior estimate of 39.61 (95% HDI: 39.03, 40.19) for the compositional patterns and a posterior estimate of 33.31 (95% HDI: 32.69, 33.93). Interestingly, the rated quality was not influenced by the length of the descriptions (*BF* = 0.01).

We also assessed how well both models captured the difficulty of communicating the different patterns, as well as participants’ quality ratings. First, we assessed whether the likelihood of each model, when fitted to the original patterns, was predictive of how communicable that pattern was. The idea behind this analysis was that, if participants were really using one of the two models to extract and compress patterns, then how well this model can compress the patterns (as measured by the likelihood given the data) should be related to how well people can communicate it. We therefore fitted a set of multi-level regression models with the previously used error measures as the dependent variables, and the log-likelihood for each pattern as estimated by both compositional and non-compositional models as the independent variables. We also included a random intercept and a random slope for each of the two models’ likelihoods, as participants might vary in their ability to redraw the described pattern and how well they are predicted by the different models. The resulting fixed effects regression coefficients (Table 2) showed the same pattern for both error measurements: there was a significant effect for the compositional but not the non-compositional log-likelihoods. Moreover, we directly compared two mixed-effects regressions solely using either the compositional or the non-compositional log-likelihoods as the independent variable. This comparison strongly favoured the compositional log-likelihoods for modeling both the absolute error (*BF >* 100) and the wavelet distance (*BF >* 100). This means that patterns that were easier to compress by the compositional model were also easier to communicate for participants. This was not true for the non-compositional model.

**Table 2.**
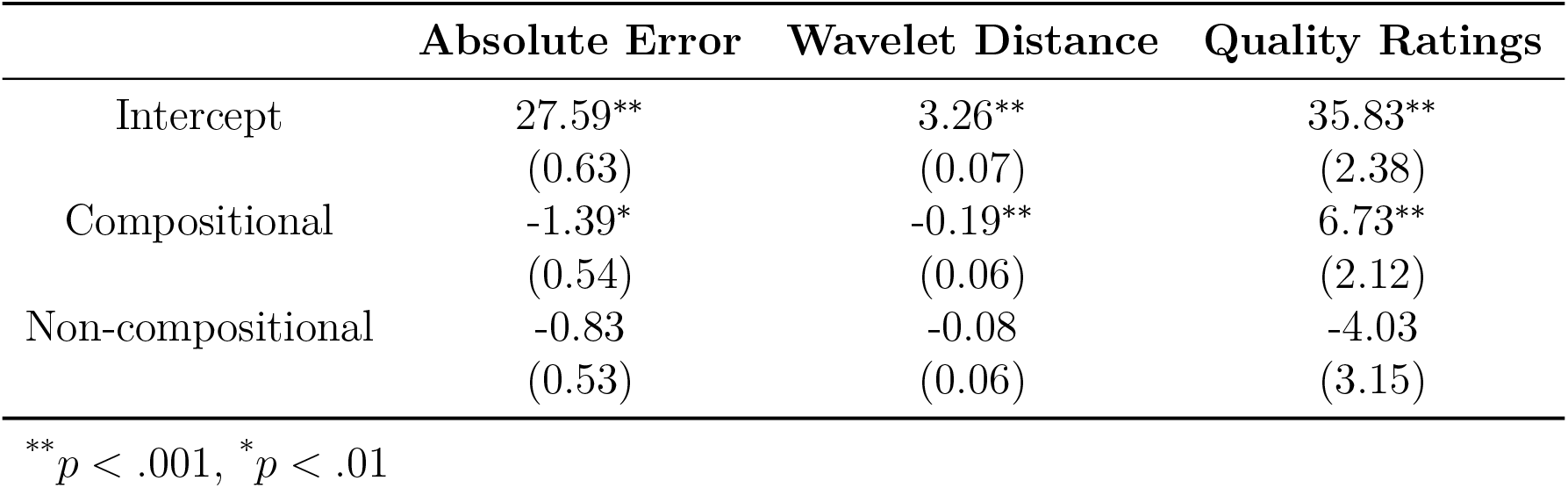
Results of regression analyses. Columns show the standardized fixed effects regression estimates for modeling the absolute error, the wavelet distance error, or participants’ quality ratings as the dependent variable. Significant effects (p < .01) are flagged by asterisks. Standard errors of the coefficients are displayed below each coefficient in brackets.

Finally, we applied the same regression approach, using the log-likelihood as the independent variable (both as a fixed and a random effect), to predict the quality ratings collected in the third part of the study. The idea behind this analysis is that if participants were indeed using one of the two models to evaluate the quality of the drawings, then they should evaluate the likelihood of the drawing to have been produced by the same generative process as the original drawing. Only the compositional model significantly predicted participant’s ratings in part 3 (Table 2 and Fig. 4c) and the direct comparison between the compositional and the non-compositional model strongly favored the compositional model (*BF >* 100). This suggests that participants assessed the quality of the drawings based on how well they could be described by similar compositions as the original patterns.

### Controlling for individual components

Given that both the compositional and the non-compositional kernel can –in the limit of infinite data– capture any function but differ in their inductive biases given finite data, we also analyzed if any individual structure (for example, periodicity or linearity) might have driven the differences between compositional and non-compositional patterns’ communicability. We therefore analyzed the differences between compositional and non-compositional patterns’ wavelet distances while controlling for how well different single component kernels described the patterns, as measured by the log-likelihoods produced by either a periodic, a linear, or a RBF kernel taken on their own. We regressed the individual components’ log-likelihoods as a fixed and a random effect onto the wavelet distances first. Additionally, we added a dummy indicating whether or not a pattern was compositional to that regression as a random effect. Afterwards, we added the same dummy variable as a fixed effect in order to assess if compositionality added something to communicability over and above the simple components. This analysis showed that adding the dummy factor improved a regression that only contained the periodic (*BF* = 20.7), the RBF (*BF* = 28.9), or the linear (*BF* = 15.6) log-likelihoods. Thus, the advantage of compositional patterns’ communicability did not solely arise from single structures, persisting even when controlling for each of the individual components of the compositional grammar.

### Controlling for pattern memorability

One concern with our current analysis is that participants saw the patterns and then had to describe them from memory. Thus, differences in the final quality could have also arisen from differences in participants’ memory capacity for different patterns. To rule out this alternative explanation, we also assessed by how much, if at all, the compositional model’s predictions captured communication quality better than just pattern memorability. We therefore ran an additional experiment in which 51 participants (37 male, mean age=31.91, SD=11.8) sequentially saw patterns for 10 seconds (just like in part 1 of our main experiment) and then had to immediately redraw it (using the same canvas setup as in part 2 of our main experiment). We let participants do this for 6 patterns in total. Three of these patterns were compositional and three were non-compositional. We then measured how well the different patterns could be remembered by calculating the wavelet differences between the original and the redrawn patterns and averaging them for each pattern individually, leading to an item-specific measure of memorability. Next, we assessed by how much our previous regressions improved by additionally entering the compositional model’s log-likelihoods as a fixed effect while controlling for the item-specific memorability score (both as a random and fixed effect) and the compositional model’s log-likelihoods as random effect. This revealed that the compositional model’s likelihood substantially improved the regression model for the absolute error (*BF* = 8.9), the wavelet distance measure (*BF* = 8.3), and the quality ratings (*BF >* 100). Thus, there are strong reasons to believe that the differences in communication qualities did not solely arise from pattern memorability.

### Relating composition-specific words to compositional descriptions

We were also interested in how specific features of participant language mapped onto specific compositions in the patterns. We therefore conducted another experiment in which we showed an additional group of participants single components of the compositional model. In this experiment, 50 participants (24 males, mean age=34.25, SD=11.96) saw 6 different patterns sequentially. Each pattern was presented to them for 10 seconds after which it disappeared and they had to describe it, exactly as in part 1 of our earlier experiments. However, this time we sampled patterns from single kernels of the compositional model. Thus, each participant had to describe 2 patterns that were sampled from a periodic kernel, 2 patterns sampled from a RBF kernel, and 2 patterns sampled from a linear kernel, presented to them in random order. We then extracted the top 10 words for each single component, i.e. the words that were more frequently used to describe patterns from a particular component compared to the other two components. The resulting words were intuitively plausible; for example, common words for periodic patterns were “peak”, “time”, and “wave”, whereas frequent words for linear patterns were “linear”, “straight”, and “steady”. We then assessed how often the extracted, composition-specific words appeared in the descriptions elicited in part 1 of our earlier experiment. Figure 5a shows how much more often the extracted words appeared in the descriptions of compositional as compared to non-compositional descriptions in our first experiment (calculated by subtracting the frequency of occurrences in the non-compositional descriptions from the frequency of occurrences in the compositional descriptions). This revealed that many of the compositional words appeared more frequently in the descriptions of compositional patterns than in the descriptions of non-compositional patterns. This can also be seen when calculating –for each set of words– the probability that at least one of the words appeared in the description (Fig. 5b). This probability was higher for compositional patterns overall (*t*(30) = 2.65, *p* = .005, *d* = 0.54, *BF* = 7.47). Moreover, both words describing periodic (*t*(30) = 4.14, *p* < .001, *d* = 0.74, *BF >* 100) and linear (*t*(30) = 3.92, *p* < .001, *d* = 0.70, *BF* = 63.3) patterns were more frequently used to described compositional than non-compositional patterns. This difference was not present for words describing RBF patterns (*t*(30) = *−*0.96, *p* = .34, *d* = 0.17, *BF* = 0.3). This is intuitive because non-compositional patterns might also contain smooth parts. Indeed, the compositional model more frequently interprets patterns sampled from the non-compositional kernel as having RBF components than linear or periodic components (cf. Schulz, Tenenbaum, et al., 2017).

**Figure 5.**
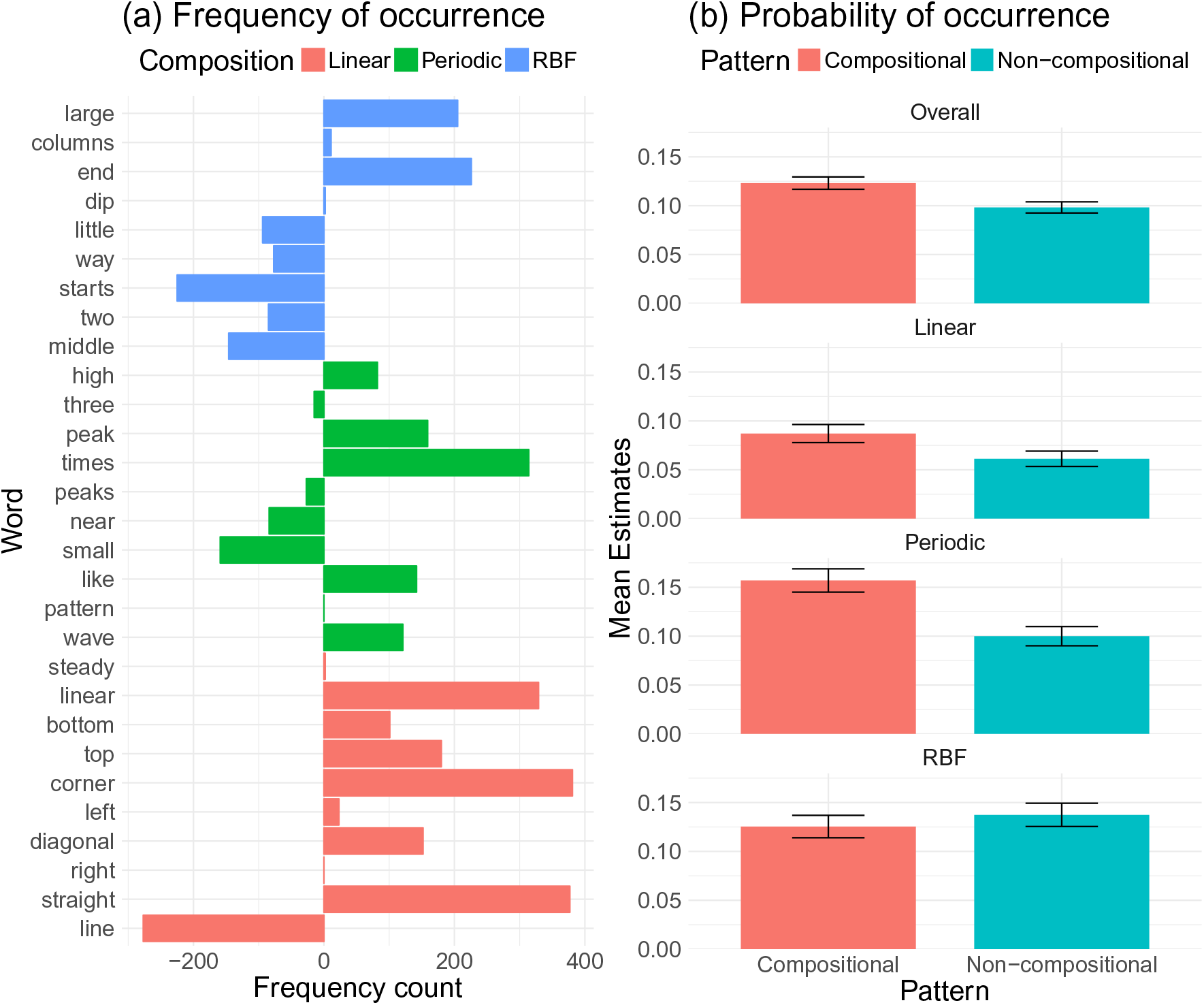
Composition-specific words. Colors indicate the type of pattern. Error bars show the standard error of the mean. a: Frequency counts of word occurrences. Words were extracted from an additional experiment asking participants to describe single patterns. Counts show how often the extracted words appeared in compositional vs. non-compositional patterns in our main experiment. For example, positive numbers show that a word extracted for a particular component appeared more frequently in the descriptions for compositional patterns than in the descriptions for non-compositional patterns. b: Probability that a word extracted from the single component descriptions would be used in a description of a compositional or non-compositional function.

Finally, we calculated for each component the probability of being present in each of the described functions. This can be approximated by dividing the summed log-likelihood of kernels containing a particular component by the sum of all log-likelihoods. We then regressed the resulting values onto a binary variable that indicated whether or not a composition-specific description was present for each description, including a random intercept over participants^3^. For example, one would expect that participants might be more likely to use RBF-specific words the more likely it actually was that a RBF component was part of the seen pattern. This showed that linear words were somewhat more likely to be used the more likely linear patterns were to be present in the data (*β* = 0.13, *z* = 2.71, *p* = .007, *BF* = 3.8, 95% HDI: .04, 0.22), and that the same was also true for RBF-specific (*β* = 0.13, *z* = 2.64, *p* = .008, *BF* = 5.2, 95% HDI: .01, 0.26) and periodic-specific words (*β* = 0.12, *z* = 2.32, *p* = .02, *BF* = 4.1, 95% HDI: .02, 0.23).

## Discussion

We investigated how people perceive and communicate patterns in a pattern communication game where one participant described a pattern and another participant used this description to redraw the pattern. Our results provide evidence that compositional patterns are more communicable, that a compositional model better captures participants’ difficulty in communicating patterns, and that participants’ quality ratings when evaluating the performance of other participants are also best captured by a compositional model. Taken together, these results suggest that there is an interface between natural language and the compositional pattern description language uncovered by our earlier work (Schulz, Tenenbaum, et al., 2017).

We are not the first to study how patterns are transmitted from one person to another. Kalish, Griffiths, and Lewandowsky (2007) let participants learn and reproduce functional patterns in an “iterated learning” paradigm. In this paradigm, participants drew functions which were then passed onto the next person, who then had to redraw them, and so forth. The results of this study showed that participants converged to linear functions with a positive slope, even if they started out from linear functions with a negative slope or just random dots. A key difference from our study is that Kalish et al. (2007) did not ask participants to generate natural language descriptions. Another difference is that in iterated learning studies, the object of interest is typically the stationary distribution, which reveals the learner’s inductive biases (Griffiths & Kalish, 2007; Kirby & Hurford, 2002). We have not attempted to simulate a Markov chain to convergence, so our study does not say anything about the stationary distribution. Here we ask whether particular pattern classes are more or less communicable. Schulz, Tenenbaum, et al. (2017) provides a systematic investigation into the nature of inductive biases in function learning, supporting the claim that these inductive biases are compositional in nature.

Our approach ties together neatly with past attempts to model compositional structure in other cognitive domains. Language (Chomsky, 1965) and object perception (Biederman, 1987) have long traditions of emphasizing compositionality. More recently, these ideas have been extended to other domains such as concept (Feldman, 2000) and rule learning (Goodman, Tenenbaum, Feldman, & Griffiths, 2008). Our results add to these attempts by linking compositional function representation to linguistic communication.

There are four important limitations of the current work, which point the way towards future research. First, we do not have a computational account of how patterns are encoded into natural language. Based on work in machine learning (Lloyd et al., 2014), one starting point is to assume that people first infer a structural description of the pattern, and then “translate” this structural description into natural language. Although the work of Lloyd et al. (2014) shows how to do this for the compositional GP model, the natural language descriptions are highly technical, and therefore a rather poor match for lay descriptions of patterns. As the word frequencies in Figures 2a-b illustrate, people seem to make use of more metaphorical language when describing compositional functions—a property not captured by the austere statistical descriptions of Lloyd and colleagues. What we need is a kind of pattern “vernacular” that maps coherently (though perhaps approximately) to the structural description.

The second limitation of our work is that we do not have a computational account of how descriptions are decoded into patterns for redrawing. One natural hypothesis is that this is essentially a reverse of the process described above: natural language descriptions are first translated into structural descriptions, which can then be plugged into the GP model to generate the mean function or sample from the posterior.

Both of these limitations might be addressed in a data-driven way by using machine learning tools to find invertible mappings from structural descriptions to natural language. In particular, we could treat this as a form of *structured output prediction*, a supervised learning problem in which the inputs and outputs are both multi-dimensional. Modern structured output prediction algorithms have developed a variety of ways to exploit the structured nature of linguistic data (e.g., Daumé, Langford, & Marcu, 2009; Tsochantaridis, Joachims, Hofmann, & Altun, 2005). These algorithms have not yet been applied to human pattern description.

The third limitation of our work is that we have investigated a fairly small set of functions. This set was chosen based on our past work (Schulz, Tenenbaum, et al., 2017) so as to minimize low-level perceptual confounds. However, further work will be required to verify that our results generalize to a broader range of functions.

The final limitation is that it is currently hard to draw a clear distinction between compositional and non-compositional patterns. Given that both the compositional and the non-compositional model can capture almost any pattern given enough data, the main differences between the two models can be derived from their predictions under a finite data regime. The two models’ inductive biases differ substantially given the number of data points we have applied here. Take as an example patterns that exhibited a linear trend. Even though the non-compositional kernel could eventually capture linear trend, it would require a large number of non-compositional parts to interpolate trends and yet would still struggle to extrapolate beyond the encountered data; this is because it lacks the required inductive biases to express trends efficiently.

## Conclusion

The idea that concepts are represented in a “language of thought” is pervasive in cognitive science (Fodor, 1975; Piantadosi, Tenenbaum, & Goodman, 2016), and we have previously shown that human function learning also appears to be governed by a structured “language” of functions (Gershman, Malmaud, & Tenenbaum, 2017; Schulz, Tenenbaum, et al., 2017). Specifically, people decompose complex patterns into compositions of simpler ones, ultimately producing a structural description of patterns that allows them to effectively perform a variety of tasks, such as extrapolation, interpolation, compression, and decision making. The results in this paper suggest that the availability of a structural description can also be used to communicate patterns in natural language. Because non-compositional functions are less effectively encoded into a structural description, they are disadvantaged in terms of accurate pattern communication. This finding provides new insight into how a language of thought might mediate translation between vision, language, and action.

## Supporting Information

### Data, descriptions and analysis code

All data, analysis script and experimental code can be found online at: https://github.com/anonymous/function_communication

All descriptions, originals and redrawn patterns can be found online at: https://anonymous.github.io/comcomppats.pdf

### Non-compositional kernel

The non-compositional kernel is based on an approximation of the spectral density of function in a Gaussian Process framework. Letting *τ* = *x* − *x*’ ∈ ℝ*^P^*, then

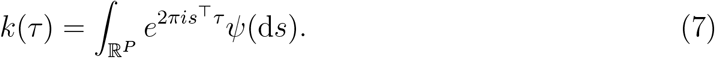

If *ψ* has a density *S*(*s*), then *S* is the spectral density of *k*; *S* and *k* are Fourier duals (Rasmussen & Williams, 2006). Thus, a spectral density over the kernel space fully defines the kernel. Furthermore, every stationary kernel can be expressed as a spectral density. Wilson and Adams (2013) showed that the spectral density can be approximated by a mixture of *Q* Gaussians, such that

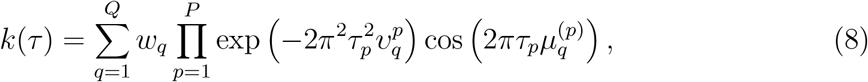

where the *q*th component has mean vector 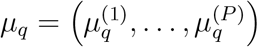 and a covariance matrix 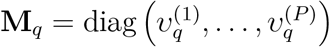.

### Statistical tests

We report all statistics using both frequentist and Bayesian tests. Frequentist tests are presented alongside their effect sizes, i.e. Cohen’s d (Cohen’s d; Cohen, 1988). Bayesian statistics are expressed as Bayes factors (BFs). A Bayes factor quantifies the likelihood of the data under the alternative hypothesis *H_A_* compared to the likelihood of the data under the null hypothesis *H*_0_. For example, a *BF* of 10 indicates that the data are 10 times more likely under *H_A_* than under *H*_0_; a *BF* of 0.1 indicates that the data are 10 times more likely under *H*_0_ than under *H_A_*. We use the “default” Bayesian *t*-test as proposed by Rouder and Morey (2012) for comparing independent groups, using a Jeffreys-Zellner-Siow prior with its scale set to 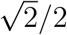. The Bayes factor for the correlation between the judges’ ratings is based on Jeffrey’s test for linear correlation as put forward by Ly, Verhagen, and Wagenmakers (2016). We approximate the Bayes factor between two different mixed-effects regressions by applying bridge sampling (Gronau et al., 2017).

### Validating the description ratings

We validated the two judges’ ratings by running an additional experiment in which 20 participants (10 females, mean age=36.2, SD=14.7, fee=$2) on MTurk sequentially rated 50 randomly sampled descriptions while also seeing the matching pattern. Participants used the same rating scale as the two judges, ranging from 1 (bad description) to 5 (very good description). The resulting mean ratings and the judges’ ratings correlated highly: *r*(29) = 0.86, *t* = 8.66, *p* < .001, *BF >* 100. Furthermore, to assess if the pre-selection had potentially biased our model comparison, we analyzed how well both models described all patterns before and after the selection. This showed no difference for either the compositional (*BF* = 0.2) or the non-compositional model (*BF* = 0.2). Moreover, both models described the selected patterns about equally well (*BF* = 0.8).

### Wavelet transform similarity measure

The discrete wavelet Haar transform performs a scale-wise decomposition of a pattern in such a way that most of the energy of the data can be represented by a few coefficients. The main idea behind this measure is to replace the original series by its wavelet approximation coefficients **a**, and then to measure the dissimilarity between the wavelet approximations. We use the R-package TSclust (Montero et al., 2014) to find the appropriate scale of the transform. We then measured the dissimilarity between two patterns *x*_1_ and *x*_2_ by the Euclidean distance at the selected scale: *d*(*x*_1_*, x*_2_) = *||***a**_1_ *−* **a**_2_*||*.

### Assessing other distance measures

We also compared compositional and non-compositional patterns using two other distance measure. The first one is the absolute distance of the actual points participants put onto the canvas and the closest points (on the x-axis) of the true patterns. This measure led to a smaller error for compositional than for non-compositional patterns (*t*(49) = 3.38, *p* = .001, *d* = 0.48, *BF* = 20.9). The second one is the absolute distance between two generalized additive models (Hastie, 2017), one fitted to participants’ drawings and one to the true underlying pattern. In contrast to the smoothing lines used in the main text, this regression was not forced to go through every point, but rather to be a more compact representation of the drawn patterns. Using this distance measure, we found the same result as before, with a smaller error for compositional than non-compositional patterns (*t*(49) = 2.72, *p* = .009, *d* = 0.38, *BF* = 4.1). We therefore conclude that compositional patterns are more communicable than non-compositional patterns, independent of the distance measure.

### Lesioned model comparison

We assessed how well the compositional model captured the difficulty of communicating different patterns (as measured by the Wavelet distance) when lesioning the model by removing kernels from the grammar. This showed that all parts of the grammar were required for good performance, since the model described errors worse without the periodic kernel (*BF* = 25.7), without the Radial Basis Function kernel (*BF >* 100) and without the linear kernel (*BF* = 48.2).

### Comparing against other smoothness-based models

We also compared the compositional kernel against multiple smoothness-only models in terms of how they captured participants’ performance. We used the Matérn class of kernel functions to encode the underlying smoothness of a function. The Matérn covariance between two points separated by *τ* distance units is

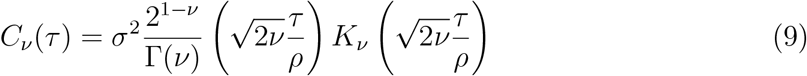

where Γ is the gamma function, *K_v_* is the modified Bessel function of the second kind, and *ρ* and *ν* are non-negative covariance parameters. For *k*_1_*_/_*_2_ the Matérn kernel is the Ornstein-Uhlenbeck kernel, which is a highly unsmooth model of functional patterns. As *k* increases, the Matérn kernel expects smoother functions. For *k → ∞* the Matérn kernel is equivalent to a Radial Basis function kernel. We compare Matérn kernels with *k* = [0.5, 1.5, 2.5, ∞] to the compositional model based on how well they predict participants’ ability to communicate patterns, measured by the wavelet distance. The compositional model predicted the communicability of patterns better than any of the smoothness-based models with all Bayes factors being larger than 100.

## Footnotes

1 Note that although the spectral kernel could a priori be captured by sums of RBF and periodic kernels, the extracted (i.e. fitted) “human kernel” reported by Wilson et al. (2015) was more similar to a mixture of a radial basis function and a linear kernel. We compare both of these types of mixture kernels to our full compositional kernel in our lesioned model comparison in the Supporting Information.

2 All descriptions can be found online: https://anonymous.github.io/comcompresps.pdf

3 We did not include a random slope over participants into this model comparison, because there was no evidence for a random slope improving model fits for the regression focusing on RBF-specific words (*BF* = 0.02), the regression focusing on linear-specific words (*BF* = 0.08) as well as the regression focusing on periodic-specific words (*BF* = 0.02).

